# Large-scale network metrics improve the classification performance of rapid-eye-movement sleep behavior disorder patients

**DOI:** 10.1101/2022.08.16.504129

**Authors:** Monica Roascio, Rosanna Turrisi, Dario Arnaldi, Francesco Famà, Pietro Mattioli, Flavio Nobili, Annalisa Barla, Gabriele Arnulfo

## Abstract

Clinical decision support systems based on machine-learning algorithms are largely applied in the context of the diagnosis of neurodegenerative diseases (NDDs). While recent models yield robust classifications in supervised two classes-problems accurately separating Parkinson’s disease (PD) from healthy control (HC) subjects, few works looked at prodromal stages of NDDs. Idiopathic Rapid-eye Movement (REM) sleep behavior disorder (iRBD) is considered a prodromal stage of PD with a high chance of phenoconversion but with heterogeneous symptoms that hinder accurate disease prediction. Machine learning (ML) based methods can be used to develop personalized trajectory models, but these require large amounts of observational points with homogenous features significantly reducing the possible imaging modalities to non-invasive and cost-effective techniques such as high-density electrophysiology (hdEEG). In this work, we aimed at quantifying the increase in accuracy and robustness of the classification model with the inclusion of network-based metrics compared to the classical Fourier-based power spectral density (PSD). We performed a series of analyses to quantify significance in cohort-wise metrics, the performance of classification tasks, and the effect of feature selection on model accuracy.

We report that amplitude correlation spectral profiles show the largest difference between iRBD and HC subjects mainly in delta and theta bands. Moreover, the inclusion of amplitude correlation and phase synchronization improves the classification performance by up to 11% compared to using PSD alone. Our results show that hdEEG features alone can be used as potential biomarkers in classification problems using iRBD data and that large-scale network metrics improve the performance of the model. This evidence suggests that large-scale brain network metrics should be considered important tools for investigating prodromal stages of NDD as they yield more information without harming the patient, allowing for constant and frequent longitudinal evaluation of patients at high risk of phenoconversion.

**Highlights:**

- Network-based features are important tools to investigate prodromal stages of PD
- Amplitude correlation shows the largest difference between two groups in 9/30 bands
- Amplitude correlation improved up to 11% the performance compared to PSD alone
- Classification robustness increases when we use both network-based EEG features
- Classifier performance worsens when PSD is added to network-based EEG features

## 1. Introduction

Nowadays, modern clinical neurology starts considering Neurodegenerative disorders (NDD) as network pathologies where connectivity between different brain areas provides more insights and predictive powers towards accurate diagnosis and disease prediction (Siegel et al., 2012). To do this, it is necessary to look for quantitative and repeatable biomarkers that characterize the brain network in neurodegenerative disease.

A good biomarker should indeed be reproducible, cost-effective, readily available, and able to serve as a disease progression marker (Miglis et al., 2021). Electroencephalography (EEG) is a non-invasive, widespread, accessible, and low-cost clinical tool used to observe dynamic changes in the neuronal electrical field and represents a gold-standard diagnostic instrument in neurology studies. Neuronal activity is hierarchically organized in different oscillations that coexist in both space and time (Buzsáki, 2006). These brain oscillations represent a mechanism for neuronal communications (Fries, 2015) where coherent field oscillations facilitate stimulus propagation within a network of brain areas. In physiological conditions, increased phase synchronization facilitates communication while abnormal increase can be predictive of several pathological conditions such as Parkinson’s disease (Roascio et al., 2021; Sunwoo et al., 2017), Alzheimer’s disease (Pusil et al., 2019), and epilepsy (Avoli, 2014; Jiruska et al., 2013).

We here focused on the Rapid eye-movement sleep Behavior Disorder (RBD) which is a parasomnia that involves violent and undesirable behaviors such as the physical reaction to dreams due to the loss of normal muscle atonia during Rapid Eye Movement (REM) sleep. People with idiopathic RBD (iRBD) have a >70% chance to develop Parkinson’s Disease (PD), Dementia with Lewy Bodies (DLB), or multiple system atrophy (Postuma et al., 2019). However, iRBD is a heterogeneous disorder and this makes accurate prediction of phenoconversion challenging.

Many studies highlighted electrophysiological changes in iRBD patients (Fantini et al., 2003; Roascio et al., 2021; Rodrigues Brazète et al., 2016; Sunwoo et al., 2017) suggesting the importance of EEG features in differentiating iRBD from healthy subjects. It is known that the “slowing down” of the power spectrum is a typical characteristic of people with iRBD compared to healthy subjects (Fantini et al., 2003; Rodrigues Brazète et al., 2016). However, the power spectrum is a frequency-wise measure of the amplitude distribution of individual cortical populations and lacks information about large-scale network interactions (Siegel et al., 2012). On the other hand, it was recently observed that phase synchronization is reduced in iRBD patients compared to healthy subjects (Sunwoo et al., 2017). Moreover, phase synchronization increases in the alpha band, while amplitude correlation decreases in the delta band with the disease progression of iRBD patients (Roascio et al., 2021).

### 1.1 Problem statement

In this study, we hypothesized that measures of the association between brain areas are likely to provide more detailed information than Fourier-based measures allowing better discrimination between RBD and healthy subjects. We thus aimed at quantifying the accuracy gained in a classification task by adopting advanced network-based EEG features in combination with more standard power-based measures.

### 1.2 State of the art

Previous studies mainly adopted clinical scores (Prashanth et al., 2016), cerebrospinal fluid (CSF) (Prashanth et al., 2016; Wang et al., 2020), magnetic resonance imaging (MRI) (Farina et al., 2020; Noor et al., 2019), single-photon emission computed tomography (SPECT) (Prashanth et al., 2014, 2016; Wang et al., 2020) as input variables to NDDs/healthy classification models. However, other studies tried to use power-based EEG metrics to investigate NDDs/healthy classification, (Bevilacqua et al., 2015; Farina et al., 2020; Lehmann et al., 2007; Maitín et al., 2020; Ruffini et al., 2016). However, few studies investigated the classification of prodromal stages of NDDs and healthy subjects by adopting EEG metrics (Dauwels et al., 2010).

In iRBD/healthy classification, the gold-standard instrument is the video-polysomnography (video-PSG) according to international criteria (ICSD 3) (Sateia, 2014). However, video-PSG requires time, resources, and highly specialized personnel. Also in this case, therefore, the clinicians looked for other criteria to support the diagnosis such as a cognitive and motor assessment battery and EEG recordings. The previous study indeed used clinical scores and gait parameters obtaining a classification accuracy of 95%, with a sensitivity of 91% (Cochen De Cock et al., 2022). Another study observed if there is useful information on the EEG spectrum to classify iRBD/healthy (Buettner et al., 2020). To do this, a Random Forest was trained using power-based features (*i.e*., power spectrum) as model input variables and obtaining an accuracy of 90% (Buettner et al., 2020). However, iRBD patients and HC subjects have different age ranges which could introduce a bias into the classification model.

Finally, a study investigated which synchrony-based EEG features better differentiate patients with mild cognitive impairment (MCI) - a prodromal stage of Alzheimer’s disease - from healthy subjects (Dauwels et al., 2010). However, to the best of our knowledge, no study has used large-scale network-based EEG features to classify people with iRBD and healthy subjects.

### 1.3 Our contribution

We investigated a cross-sectional cohort of 105 subjects: 59 people with iRBD and 46 healthy subjects. We recorded high-density EEG signals during relaxed wakefulness, and we investigated if large-scale EEG features (*i.e*., phase synchronization and amplitude correlation) are more discriminative than the power spectrum to differentiate iRBD from healthy subjects. We then used the EEG-based features as input variables for a machine learning (ML) model with sparsity, to perform iRBD/healthy classification. The sparsity-based regularization allowed us to identify the EEG feature(s) that are most important for training the classification model. Finally, we used this information to train a regression model without sparsity for evaluating the classification performance when using a subset of variables - chosen based on the previous model.

## 2. Materials and methods

### 2.1 Materials

#### 2.1.1 Sample characteristics

In collaboration with the sleep lab of the Clinical Neurology, University of Genoa, IRCCS Policlinico San Martino, we recruited a total of 110 subjects (Table 1) including 62 iRBD patients (9 female; mean age 69.61±6.98 years at the first clinical evaluation) and 48 healthy subjects (23 female, 70.25±10.26). After 23.82±18.13 months, 19 iRBD converted to α-synucleinopathy (PD=8; DLB=11).

**Table 1:** Main clinical and demographic data of idiopathic RBD patients and HC subjects. **Legend**. Idiopathic Rapid eye-movement sleep Behavior Disorder – iRBD, Healthy control subjects - Controls, Mini-Mental State Examination - MMSE, Movement Disorder Society-sponsored revision of the unified Parkinson’s disease rating scale, motor section - MDS-UPDRS-III, Parkinson’s Disease Sleep Scale version 2 - PDSS-2, Beck Depression Index II - BDI-II.

|  | iRBD | Controls |
| --- | --- | --- |
| Age (years) | $69.61 \pm 6.98$ | $70.25 \pm 10.26$ |
| Sex, F (%) | 9 (14.5%) | 23 (48%) |
| MMSE | $28.04 \pm 2.19$ | $28.55 \pm 2.29$ |
| MDS-UPDRS-III | $1.71 \pm 3.21$ | 0 |
| BDI-II | $11.07 \pm 7.86$ | n.a. |
| PDSS-2 | $14.16 \pm 8.01$ | n.a. |

#### 2.1.2 Inclusion criteria and ethical committee approval

We diagnosed the iRBD according to international criteria (ICSD 3) (Sateia, 2014) and we confirmed the diagnosis with overnight video polysomnography. All iRBD patients have been subjected to brain Magnetic Resonance Imaging (MRI) or Computed Tomography (CT), to rule out brain diseases such as tumors or lesions. We did not use the presence of white matter as exclusion criteria if the Wahlund scale was not >1 for all brain regions (Wahlund et al., 2001).

All subjects underwent general examinations to rule out other neurological and psychiatric disorders. In particular, the clinical evaluation for iRBD subjects included: i) the Mini-Mental State Examination (MMSE) as a measure of global cognitive function; ii) the Movement Disorder Society-sponsored revision of the unified Parkinson’s disease rating scale, motor section (MDS-UPDRS-III) to investigate the presence of parkinsonism; iii) clinical interviews and questionnaires for activities of daily living (ADL) and instrumental ADL to exclude dementia; iv) the Beck depression inventory II (BDI-II) to evaluate depressive symptoms; v) the Italian version of Parkinson Disease Sleep Scale version 2 (PDSS-2) to measure the sleep disorders (Arnaldi et al., 2016). For HC subjects, we evaluated the MMSE, and we set the MDS-UPDRS-III equal to zero because healthy subjects were free of Parkinsonian signs.

The study was conducted under the declaration of Helsinki, and all participants gave informed consent before entering the study, which was approved by the local ethics committee.

#### 2.1.3 Data collection and availability

To minimize drowsiness, prevent sleep, and preserve a high signal quality, we recorded the EEG in the morning, and we monitored the session to maintain a constant level of vigilance of the patient. We thus used a high-density (64 channels) cap of the Galileo system (EBNeuro, Florence, IT) to acquire band-passed (0.3 – 100 Hz) signals during relaxed wakefulness, using a sampling rate equal to 512 Hz. We adopted the 10-10 International System to put on the electrodes on the cap. We chose Fpz and Oz as the reference electrode and ground, respectively. To check eye movements, we synchronically acquired the horizontal electrooculogram with the same recording parameters of EEG. Finally, we checked that the electrode impedance was below 5kOhm.

### 2.2 Methods

#### 2.2.1 Data preparation

First, we pre-processed the high-density EEG data using Brainstorm (Tadel et al., 2011), a MATLAB R2021a toolbox. We applied a zero-phase infinite response notch filter (order 2) to remove the power line noise (50Hz). We thus removed all channels (mean 1.90±2.29; range: min 0, max 8) and windows showing artefactual activity using both Independent Component Analysis (ICA) and visual inspection. For all subjects, we discarded channels A1, A2, and POZ due to the presence of artifacts in more than 90% of the recording. We thus applied a band-pass finite impulse response filter (1-80 Hz, Kaiser window, order 3714). Later, we interpolated the bad channels using spline interpolation (kernel size: 4 cm). Finally, we transformed the referenced EEG to Scalp Current Densities (SCD) (Perrin et al., 1989) to all clean sensors with a spline method (lambda 0.00001, stiffness 4).

After the pre-processing, five subjects were excluded due to excessive artefactual activity, which left less than 3 minutes of pruned eyes-closed resting-state data. The final population size for this study was 105 subjects, including 59 iRBD patients (9 female; mean age 69.28±6.98 years at the first clinical assessment) and 46 healthy subjects (22 female, 70.5±10.32).

#### 2.2.2 Feature extraction

For each SCD time-series, we extracted power spectral profile, phase-synchronization, and amplitude correlation.

First, we computed the Power Spectral Density (PSD) for all subjects using the Welch algorithm with 1 Hz resolution as a standard reference to compare with current literature.

We then quantified the large-sale brain network alterations by evaluating the weighted Phase Lag Index (wPLI) (Vinck et al., 2011), which is a measure of synchronization of the phase of two signals (*i.e*., phase synchronization), and orthogonalized Correlation Coefficient (oCC) (Hipp et al., 2012) that estimates the correlation of the envelopes of two signals (*i.e*., amplitude correlation). Specifically, we conducted a time-frequency decomposition using 30 narrow-band Morlet wavelets in a logarithmic space between 2.1 and 75 Hz with 5 cycles. For each Morlet wavelet, we computed the wPLI and the oCC.

The wPLI is computed as (Vinck et al., 2011):

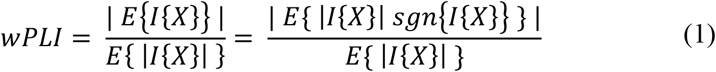

The oCC is computed as the Pearson correlation coefficient between two orthogonalized time series (Hipp et al., 2012):

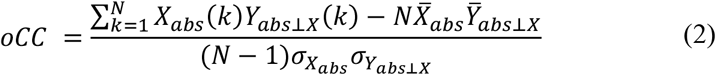

Both wPLI and oCC range between 0-1 with 1 corresponding to the presence of phase-synchronization or amplitude correlation.

We thus chose these two EEG features because they are measures insensitive to volume conduction (Hipp et al., 2012; Vinck et al., 2011), which would inflate phase synchronization and amplitude correlation analyses of sensor EEG data (Palva et al., 2018; Vinck et al., 2011).

We thus obtained a total of 3 features to classify RBD and healthy subjects: (i) power spectrum profile (2-40 Hz); (ii) phase synchronization (2.1–70.2 Hz); and (iii) amplitude correlation (2.1-70.2Hz).

Please note that in this study, we did not include clinical features in the model input, although this could potentially improve the model performance. Indeed, the main goal of the study is to investigate the discrimination power of the EEG features and determine which one is more informative for the classification task.

Further, we did not include sex information in the input classifier as our population is sex imbalanced (N=50 males with iRBD and N=24 healthy males) and, again, this may represent a model bias. This imbalance is not specific to our dataset, but it is known that there is a male predominance in iRBD patients (Arnaldi et al., 2021; Postuma et al., 2019). Nonetheless, we carried out an explorative analysis where we also used age and sex as input variables together with EEG features to classify patients with iRBD and healthy subjects (Figure S2).

#### 2.2.3 Statistical Analysis

We conducted a statistical analysis to investigate differences between people with RBD and healthy subjects. We computed a Kruskal-Wallis test (non-parametric ANOVA) (Kruskal & Wallis, 1952) to observe a statistical difference between RBD/healthy subjects in the clinical scores, power spectrum, wPLI, and oCC across frequency. Later, we performed a multiple comparison correction using the Benjamini-Hochberg (BH) method (Benjamini & Hochberg, 1995).

#### 2.2.4 Machine Learning methods for classification

We performed iRBD/healthy subject classification based on a machine learning model. A crucial remark is that we only have access to a small amount of data to train our model. This might cause overfitting, a very well-known issue in ML, which occurs when the classifier learns noise and random fluctuations in the training data and does not generalize to the test data (*i.e*., unseen data). The main causes of overfitting are indeed a small number N of samples in training or the high complexity of the model (*e.g*., large number P of input variables).

A strategy to prevent overfitting due to the high complexity of the model is the selection of a subset of variables to use as input to the model, so that P≪N. We thus adopted the Least Absolute Selection and Shrinkage Operator (LASSO), which uses the ℓ_1 norm for regularization and performs an automatic variable selection. Furthermore, we carried out additional analysis to investigate if directly using the selected variables based on LASSO as model input is advantageous in iRBD/healthy classification and how the performance changes with the input dimension P. To do that, we first derived from the LASSO results in a strategy to select subgroups of variables that may be more informative for the classification (see *Experimental design* 2.2.5 section for details) and we then used these as input to train a classification model without sparsity.

#### 2.2.5 Experimental design

The experiments we carried out can be subdivided into two phases (Figure 1a).

**Figure 1:**
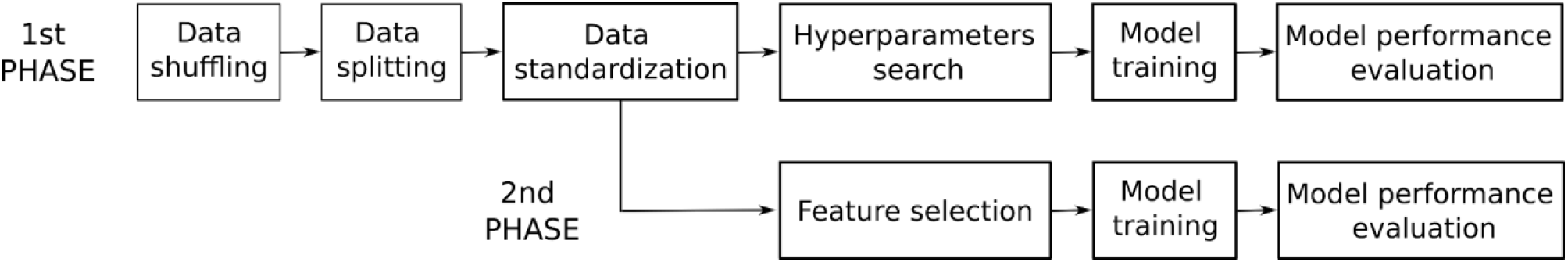
Experimental design. Summary of the two phases of the experiments. In the first phase, we carried out three different shuffling of the data. For each shuffle, we split the data into 5 different folds using the stratified 5-folds cross-validation and then we standardized the data. Later, we carried out a hyperparameters search, and then, we trained a model with the best parameters. In the second phase, we trained a regression model without sparsity using as input variables the variables selected in the previous phase based on LASSO, and then we evaluated the model performance. **Legend**. Least Absolute Selection and Shrinkage Operator – LASSO.

In the first phase, we adopted LASSO to simultaneously perform variable selection and binary classification. Specifically, here we used different input variables containing: (i) a single EEG-based feature; (ii) a pair of EEG-based features; or (iii) all EEG-based features for all 105 people involved. We then performed three different shuffling of the population data, and, for each shuffling, we split the dataset in learning and testing by stratified 5-fold cross-validation (outer-CV) (Farina et al., 2020; Kiiski et al., 2018). For each fold, we thus applied data standardization by computing the z-score on the learning set and applying the same transformation to the test set. We then further split the learning set into training and validation sets, named inner-CV (Figure 1b), for tuning the model hyperparameters. Hence, we looked for the best ℓ_1regularization parameter α (among 0.001, 0.01, 0.1, 1) by choosing the one that produced the lowest prediction error on the validation set, and we trained a Lasso model on the learning set (Figure S1). Subsequently, we evaluated the model performance in terms of accuracy, precision, recall, and f1-score on the unseen test set. The accuracy score indicates the percentage of labels predicted correctly. The precision score is defined as the ability of a classifier to not mislabel a sample (TP/TP+FP where TP and FP are true and false positive, respectively). The recall, or sensitivity, score is the ability of a classifier to find all the positive samples (TP/TP+FN where FN is the false negative). The f1-score is a weighted harmonic mean of precision and recall. We, finally, computed the mean and the standard deviation of these scores across folds. We repeated this procedure three times, shuffling the original data each time, to test the robustness of the model to the dataset split.

In the second phase, we carried out further analysis for evaluating the classification performance when using a subset of variables. To do that, we first considered the shuffling (N=3) and the folds (M=5) of the previous phase as 15 (M x N) different folds. For each one of these folds, we looked at which variables have been selected based on LASSO, and then, we counted how many folds (*i.e*., how many times) a variable was selected. We thus created different variable subsets considering the variables selected in at least one-fold up to those selected in all folds and we used them as input for a sparsity-free model.

For each input subset, we again performed three different shuffling of the data - the same as used for LASSO - and the outer-CV to provide a more robust evaluation of the model. We thus trained the model without sparsity on the learning set and then, we evaluated the model on the test set computing the accuracy, precision, recall, and f1-score. We finally calculated the averaged performance and the standard deviation across folds and shuffling. Please note that in this phase, we do not perform any inner cross-validation as we do not have a hyperparameters search.

## 3. Results

### 3.1 EEG features change in iRBD patients

We first wanted to characterize the differences between iRBD and HC populations by performing statistical hypothesis tests of single EEG features across frequencies. Power spectral profile shows a slowdown of the prominent alpha peak towards delta/theta band in iRBD patients compared to healthy subjects. This difference does not reach statistical significance after correction for multiple comparisons (*p*>0.05 - Kruskal-Wallis test and BH correction) (Figure 2a). Phase synchronization is weaker in iRBD patients than in healthy subjects in delta, theta, and alpha (8-13 Hz), but with no significant evidence (*p*>0.05 - Kruskal-Wallis test and BH correction) (Figure 2b). Finally, amplitude correlation is stronger (*p*<0.05 - Kruskal-Wallis test and BH correction) in iRBD patients than in healthy subjects in delta (2-4Hz), theta (5-7Hz), high-beta (20-30Hz), and gamma (>30Hz) bands (Figure 2c). These differences are not significant after BH correction *(p>* 0.05 - Kruskal-Wallis test and BH correction).

**Figure 2:**
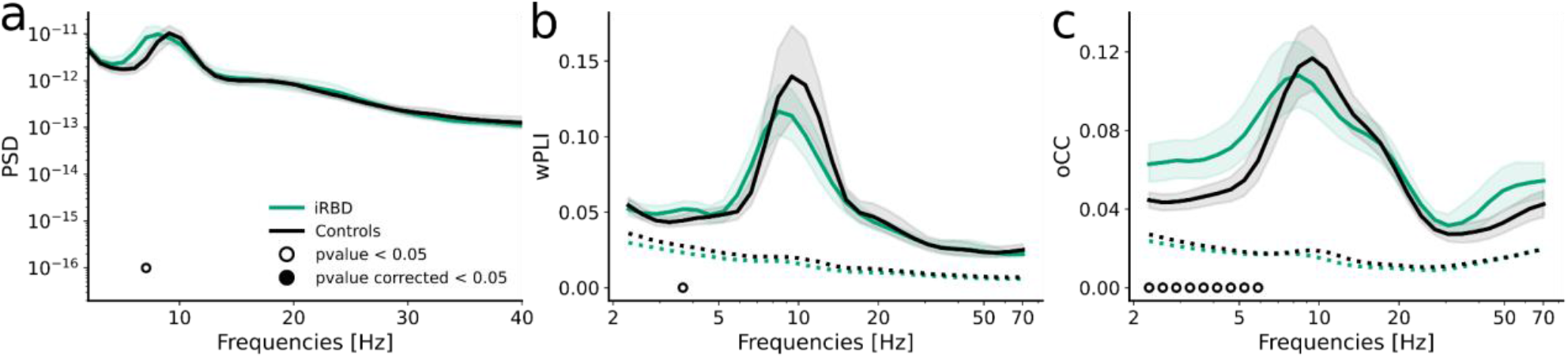
EEG features change in iRBD patients. Group-level averaged (continuous lines) and single-subject (dotted lines) of (a) PSD, (b) wPLI, and (c) oCC for 2-classes: RBD (green) and controls (black). Shaded areas represent confidence intervals at 5% around population mean (bootstrap, N = 1000). Dashed lines in (b) and (c) represent surrogate averages. Empty black circles show a significant difference between people with RBD and healthy subjects (Kruskal Wallis test). Filled black circles show a significant difference between the two population after the BH correction. **Legend**. Power spectral density – PSD, weighted Phase Lag Index – wPLI, orthogonalized Correlation Coefficient – oCC, idiopathic Rapid eye-movement sleep Behavior Disorder – iRBD, Healthy control subjects – Controls.

Despite the lack of significance after correction per multiple comparison, these results suggest that the amplitude correlation spectral profile yields major differences between iRBD, and HC compared to power-based spectral metrics.

### 3.2 Large-scale EEG features improve classifier robustness and performance

We then wanted to quantify the improvement of classification accuracy when using a combination of large-scale network metrics compared to purely power-based metrics. We quantified the classifier performance in terms of f1-score, accuracy, precision, and recall. We showed stronger robustness (*i.e*., less variability across folds and shuffling) of the classifier when using simultaneously large-scale EEG features than other feature groups (Figure 3a). Moreover, the model overperforms the PSD-based classification when using only oCC (f1-score: 61%) or both large-scale EEG features (f1-score: 62%) (Figure 3b-e). We found that the PSD-based model only reaches the chance level. Furthermore, the performance worsened when we added PSD to the large-scale EEG features compared to the performance obtained using the large-scale EEG features alone or together.

**Figure 3:**
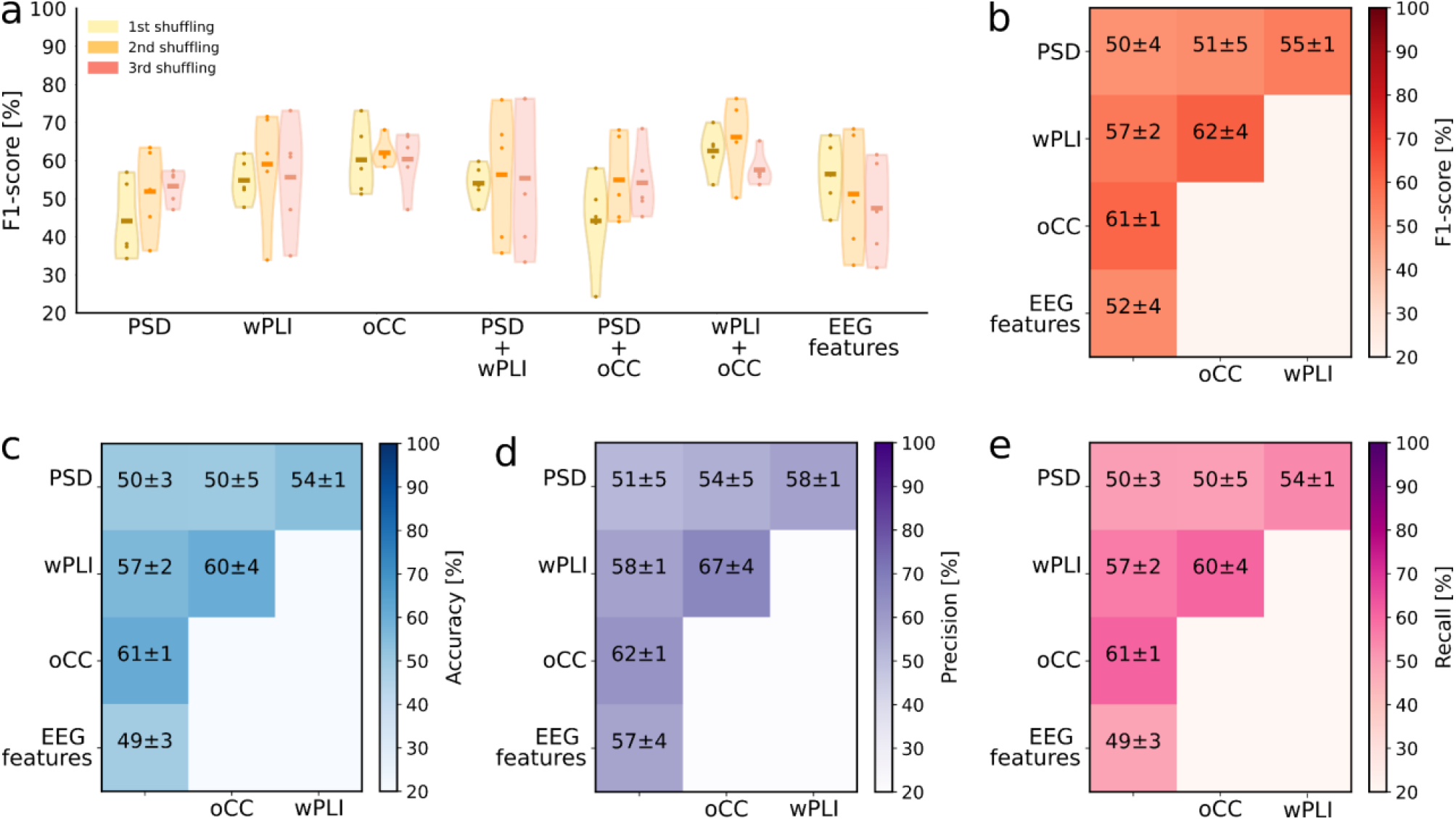
Large-scale EEG features improve the robustness and performance of the classifier. (a) Violin plots of F1-score of classification model with 7 different inputs of variables as reported on the x-axis. For each model, we used three different shuffling (yellow, orange, and light red, respectively) to shuffle data before the split in learning and testing sets. (b-e) Averaged and standard deviation across shuffling of (b) f1-score (red), (c) accuracy (blue), (d) precision (purple), and (e) recall (pink). **Legend**. Power spectral density – PSD, weighted Phase Lag Index – wPLI, orthogonalized Correlation Coefficient – oCC, Electroencephalographic – EEG.

These results are thus in line with our previous statistical findings (Figure 2) for which the large-scale EEG features, and in particular amplitude correlation profiles might be more informative in the iRBD/healthy discrimination.

Finally, we carried out a further analysis to investigate the classifier performance by adding age and sex as additional features. As iRBD is primarily a male disease, we found that the performance of the classifier improves (Figure S2) suggesting that sex imbalance in our population may represent a bias in the model.

### 3.3 Performance increased by adopting less than 50% of the features as input variables

Underpowered classification studies might benefit from feature selection. Here we investigated the group of EEG features that provided the best performances.

Firstly, we found that excessive feature selection was disadvantageous when we used wPLI alone or wPLI and oCC together. Indeed, the performance of the classifier worsened - below the chance level - when we used less than 15% of the variables as model inputs (Figure 4 and S3). In the same way, the performance worsened when we trained the model using the whole spectrum of the EEG feature(s). In contrast, the performance improved when we used less than 50% of the variables as input for the binary classification model (Figure 4).

**Figure 4:**
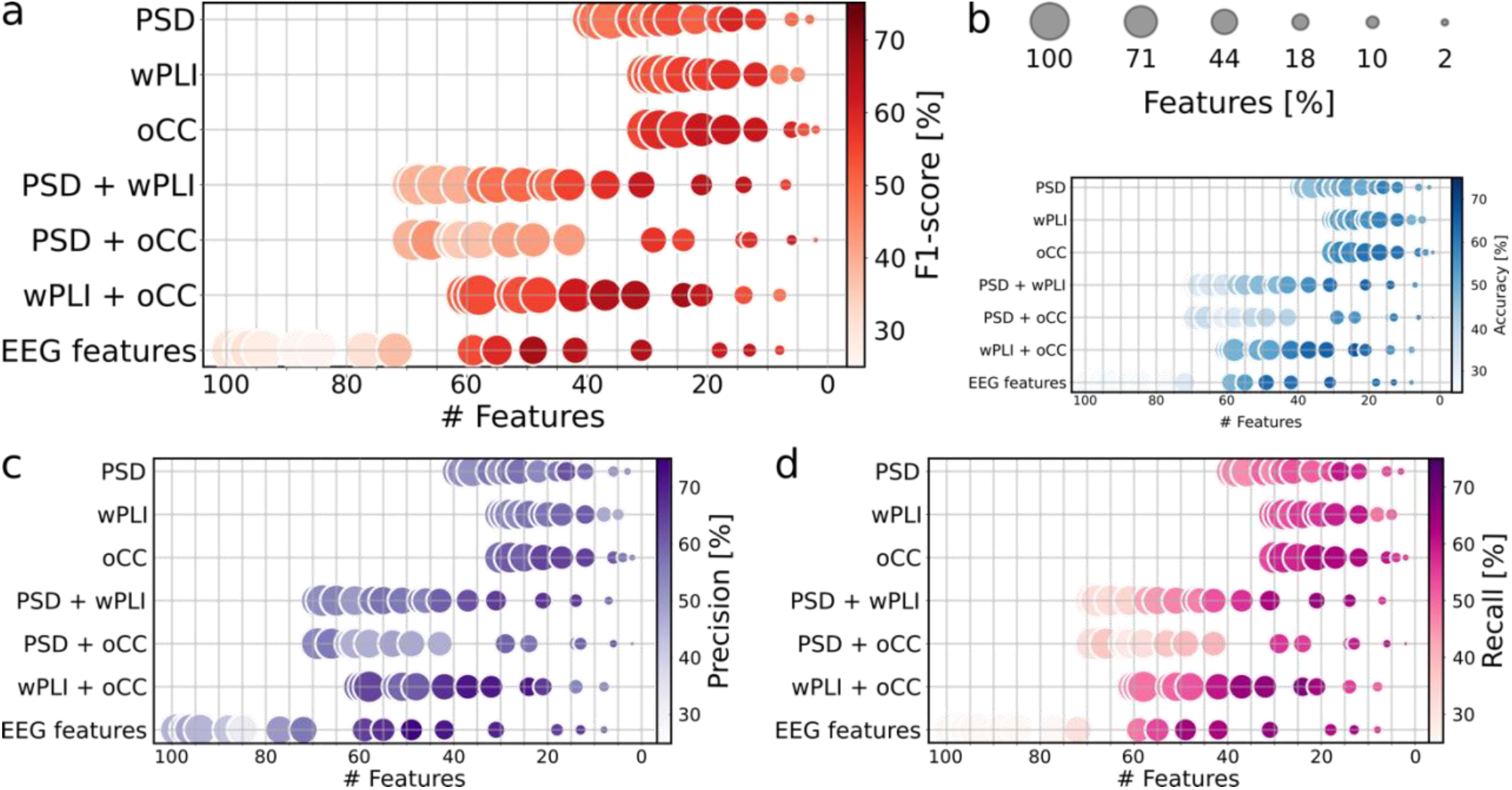
Performance increased by adopting less than 50% of the features as input variables. Averaged and standard deviation of (a) f1-score (red), (b) accuracy (blue), (c) precision (purple), and (d) recall (pink) across shuffling and split for different input variables. The first bubble for each group of features shows the performance of the classifier without variable selection. Other bubbles show the classifier performance with features selection (the number of selected variables is shown on the x-axis). The bubble size shows the percentage of selected variables, while the bubble color shows the performance of the model. On the y-axis, it was shown the group of features that are used as input variables to carry out the RBD/healthy classification. **Legend**. Power spectral density – PSD, weighted Phase Lag Index – wPLI, orthogonalized Correlation Coefficient – oCC, idiopathic Rapid eye-movement sleep Behavior Disorder – iRBD.

We found several groups of features with similar improved performances (Figure 4, S3). In particular, the oCC alone showed an f1-score of 63% when selecting 17 of its frequency points (Figure 4). The wPLI and oCC together showed the best performance (f1-score: 69%) when we selected 40% of the variables in these two groups (Figure 4). We obtained a similar performance when we selected 49%% of the variables considering all EEG features together (Figure 4). Finally, we found (Figure 4) a worsening of the performance when we trained the model using amplitude correlation or phase synchronization and power spectrum (f1-score: 65% PSD + wPLI; f1-score: 62% PSD + oCC). We observed large variability in the percentages of the selected feature number due to the different sizes of the original training set. However, we showed that the optimal number of features is between 10 and 40 (Figure 4).

These results suggest that the whole frequency spectrum of the EEG features is unnecessary for iRBD/healthy classification, agreeing that the idea to use LASSO in the first phase of the experimental design is appropriate. However, we did not find a specific subset of EEG features that performs the classification much better than others.

## 4. Discussion

To predict phenoconversion in neurodegenerative diseases, ML-based approaches would require a plethora of stable biomarkers reflecting disease progression that can be homogeneously collected across time

To be effective and widely adopted in clinical practice, these biomarkers need to be non-invasive, widely accessible across different centers, and accurately separate between healthy and pathological conditions even at prodromal stages.

In the last decades, the researchers mainly used clinical scores (Prashanth et al., 2016), cerebrospinal fluid (CSF) (Prashanth et al., 2016; Wang et al., 2020), or features extracted by imaging techniques (*e.g*., MRI and SPECT) (Farina et al., 2020; Noor et al., 2019; Prashanth et al., 2014, 2016; Wang et al., 2020) as input variables to the classification models of different NDDs including Alzheimer or Parkinson’s disease and their respective main prodromal stages as Mild Cognitive Impairment and RBD, respectively.

However, CSF, PET, and SPECT are diagnostic techniques that are very expensive, invasive, and not always accessible across different centers. In this work we specifically considered EEG-only features for its technical advantages and we selected phase synchronization and amplitude correlations as EEG-derived features in contrast to classical power spectral analyses.

Previous studies highlighted significant electrophysiological changes in iRBD patients (Fantini et al., 2003; Roascio et al., 2021; Rodrigues Brazète et al., 2016; Sunwoo et al., 2017) suggesting the importance of EEG-based features in differentiating iRBD from healthy subjects. These works reported a slowing of the alpha rhythm (Fantini et al., 2003; Rodrigues Brazète et al., 2016) and alterations in phase synchronization (Roascio et al., 2021; Sunwoo et al., 2017) and amplitude correlations (Roascio et al., 2021). Several lines of evidence suggest that NDDs are characterized by network deficiencies, and that the investigation of the cross-talk between different brain areas provides a better understanding of the diseases and their progression (Pusil et al., 2019; Roascio et al., 2021; Sunwoo et al., 2017).

We hypothesized that network-based EEG features more reliably capture differences between patients and healthy controls, and this would be reflected by an improvement in the robustness and the accuracy in a classification model.

We quantified the gain in classification accuracy of patients with idiopathic REM sleep behavior disorder from age-matched healthy controls using phase synchronization and amplitude correlation profiles. Our analyses suggest that: (1) amplitude correlations yield the largest difference around alpha band, (2) accuracy increases when using a combination of network-based features compared to more classical ones derived from Fourier spectrum, (4) the inclusion of power spectrum features decreases model accuracy, and finally that (3) strong feature selection is not beneficial when using EEG-derived metrics.

Our results provide evidence that (1) EEG-based features can be used in a classification model of RBD patients, and (2) that phase synchronization and amplitude correlation should be considered as important features in diagnosis support systems as they capture subtle changes of the progressive pathologies even in absence of overt symptoms.

We acknowledge that there are some limitations to this study. First, the small number of subjects involved limits the generalizability of our observations. However, this study aims to demonstrate that network-based metrics carry more information about the ongoing pathology by observing an increase of the accuracy in a classification model. Second, the sex could introduce a bias in our classification model. As discussed in the Feature Extraction section, the sex imbalance between healthy subjects and iRBD patients is not specific to our dataset but it is known that there is a male predominance in iRBD patients (Arnaldi et al., 2021; Postuma et al., 2019). Nonetheless, we carried out an explorative analysis where we also used age and sex as input variables together with EEG features to classify patients with iRBD and healthy subjects (see Figure S2). As expected, we found a performance improvement when using age and sex combined with EEG feature(s).

## 5. Conclusion

This study, for the first time, investigates if the network-based features improve the robustness and the classification performance in a cross-sectional cohort of iRBD patients and healthy subjects. Our results suggest that the power spectrum alone is not discriminative enough to perform an accurate iRBD/healthy classification reaching only the chance-level. In contrast, phase synchronization and amplitude correlation increased classifier performance compared to PSD alone, and classifier robustness improved when we simultaneously used both as input for the model. These findings suggest that network-based EEG features are more discriminative than power-based EEG features to improve the robustness and the performance of a classification model. We speculate that to accurately predict phenoconversion of RBD patients using ML-based approaches EEG derived features need to be included among the observed variables, in particular phase synchronization and amplitude correlations.

## Data/code availability statement

The data used in this work can be available upon a reasonable request due to privacy issues of clinical data.

The Python code is available at: https://github.com/MonicaRoascio/ClassificationRBD.git

## CRediT author statement

**Monica Roascio**: Conceptualization, Formal Analysis, Methodology, Writing – Original Draft, Writing-Reviewing and Editing, Visualization. **Rosanna Turrisi**: Conceptualization, Methodology, Writing-Reviewing and Editing. **Dario Arnaldi**: Investigation, Resources, Writing-Reviewing and Editing. **Francesco Famà**: Investigation, Writing-Reviewing and Editing. **Pietro Mattioli**: Investigation, Writing-Reviewing and Editing. **Flavio Nobili**: Resources, Writing-Reviewing and Editing. **Annalisa Barla**: Conceptualization, Writing-Reviewing and Editing, Supervision. **Gabriele Arnulfo**: Conceptualization, Writing-Reviewing and Editing, Supervision.

## Acknowledgements

This work was developed within the framework of the DINOGMI Department of Excellence of MIUR 2018–2022 (legge 232 del 2016).

This work was supported by EU H2020 Virtual Brain Cloud (grant number: 826421); and grant from Italian Ministry of Health - Italian Neuroscience network (RIN).

Rosanna Turrisi was supported by a research fellowship funded by the DECIPHER-ASL – Bando PRIN 2017 grant (2017SNW5MB - Ministry of University and Research, Italy).

Dario Arnaldi discloses lecture and consulting fees from Fidia, Jazz, Bioprojet and Lundbeck.

