## Supplementary materials for "Large-scale network metrics improve the classification performance of rapid-eye-movement sleep behavior disorder patients"

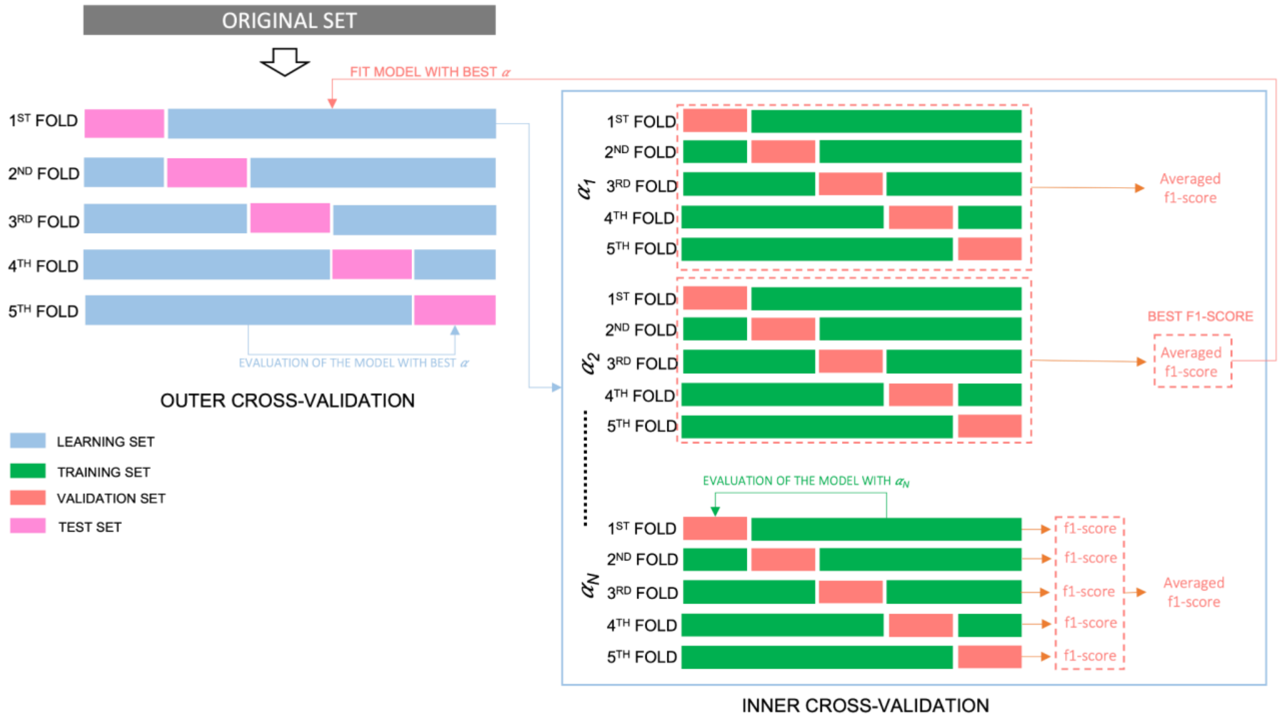


*Figure S1: **Nested cross-validation.** In the first phase of the experimental design, we used both outer and inner cross-validation (CV). The outer-CV to split the original dataset in k different folds, and the inner-CV to tune the model hyperparameters to search the best regularization parameters for LASSO. Please note that we performed the outer cross-validation, but not the inner cross-validation in the second phase of the experimental design.*

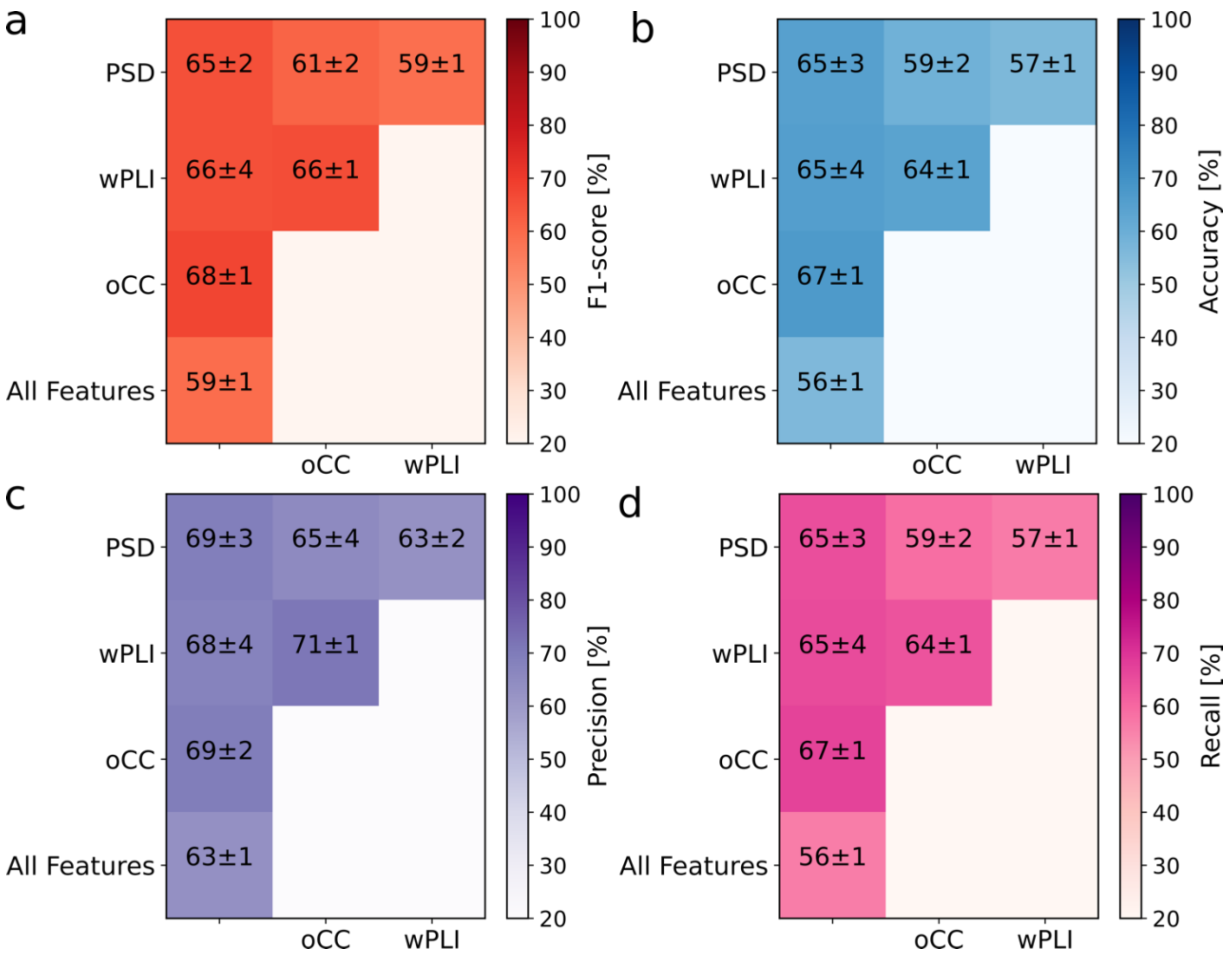


*Figure S2:Averaged classifier performance and standard deviation across shuffling of f1-score (a), accuracy ba), precision (c), and recall (d) using as inputs to classifier both demographics features (age and sex) and EEG features.* ***Legend****. Power spectral density – PSD, weighted Phase Lag Index – wPLI, orthogonalized Correlation Coefficient – oCC, Electroencephalographic – EEG.*

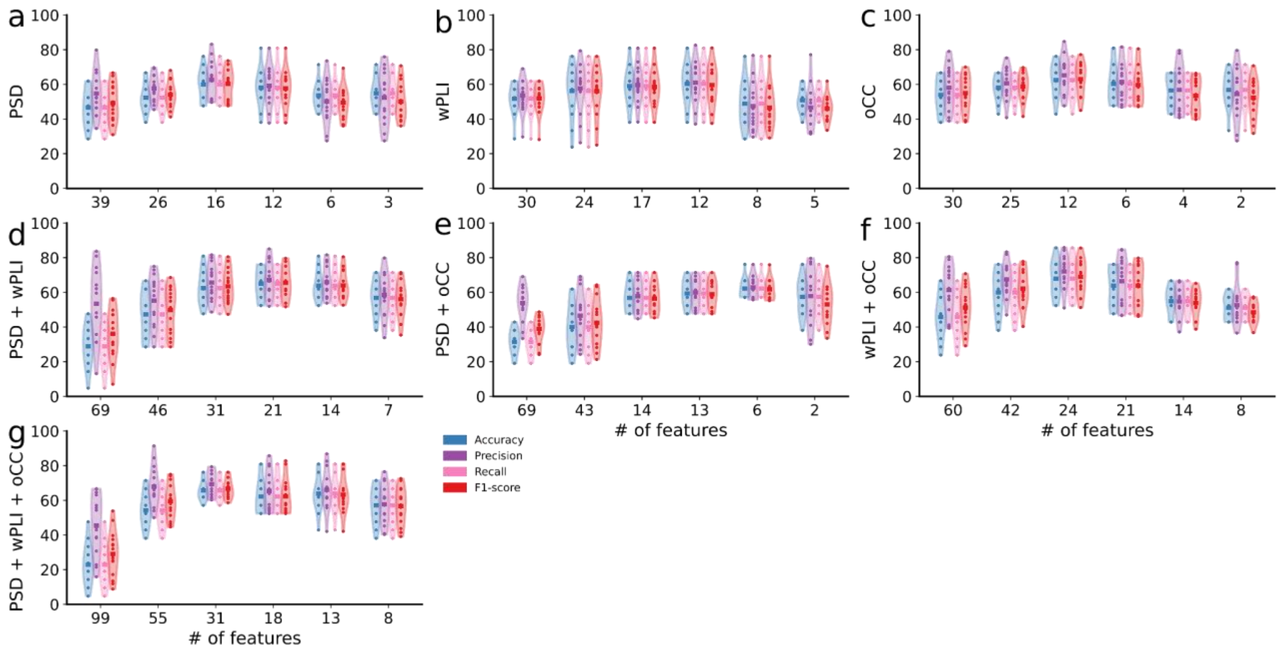


*Figure S3: Violin plots of f1-score (red), accuracy (blue), precision (purple), and recall (pink) of classification model with 7 different input variables: (a) PSD, (b) wPLI, (c) oCC, (d) PSD + wPLI, (e) PSD + oCC, (f) wPLI + oCC ,and (g) PSD + wPLI + oCC. Legend. Power spectral density – PSD, weighted Phase Lag Index – wPLI, orthogonalized Correlation Coefficient – oCC.*
